# Morphotype-Resolved Characterization of Microalgal Communities in a Nutrient Recovery Process with ARTiMiS Flow Imaging Microscopy

**DOI:** 10.1101/2024.12.16.628756

**Authors:** Benjamin Gincley, Farhan Khan, Md Mahbubul Alam, Elaine Hartnett, Ga-Yeong Kim, Hannah R. Molitor, Autumn Fisher, Ian Bradley, Jeremy Guest, Ameet J. Pinto

## Abstract

Microalgae-driven nutrient recovery represents a promising technology to reduce effluent phosphorus while simultaneously generating biomass that can be valorized to offset treatment costs. As full-scale processes come online, system parameters including biomass composition must be carefully monitored to optimize performance and prevent culture crashes. In this study, flow imaging microscopy (FIM) was leveraged to characterize microalgal community composition in near real-time at a full-scale municipal wastewater treatment plant (WWTP) in Wisconsin, USA, and population and morphotype dynamics were examined to identify relationships between water chemistry, biomass composition, and system performance. Two FIM technologies, FlowCam and ARTiMiS, were evaluated as monitoring tools. ARTiMiS provided a more accurate estimate of total system biomass, and estimates derived from particle area as a proxy for biovolume yielded better approximations than particle counts. Deep learning classification models trained on annotated image libraries demonstrated equivalent performance between FlowCam and ARTiMiS, and convolutional neural network (CNN) classifiers proved significantly more accurate when compared to feature table-based deep neural network (DNN) models. Across a two-year study period, *Scenedesmus* spp. appeared most important for phosphorus removal, which were negatively associated with elevated temperatures and nitrite/nitrate concentrations. *Chlorella* and *Monoraphidium* also played an important role in system performance. For both *Scenedesmus* and *Chlorella*, smaller morphological types were more often associated with high system performance, whereas larger morphotypes implied a stress response correlating with poor phosphorus recovery rates. These results demonstrate the potential of FIM as a critical technology for high-resolution characterization of industrial microalgal processes.

**Graphical Abstract:** 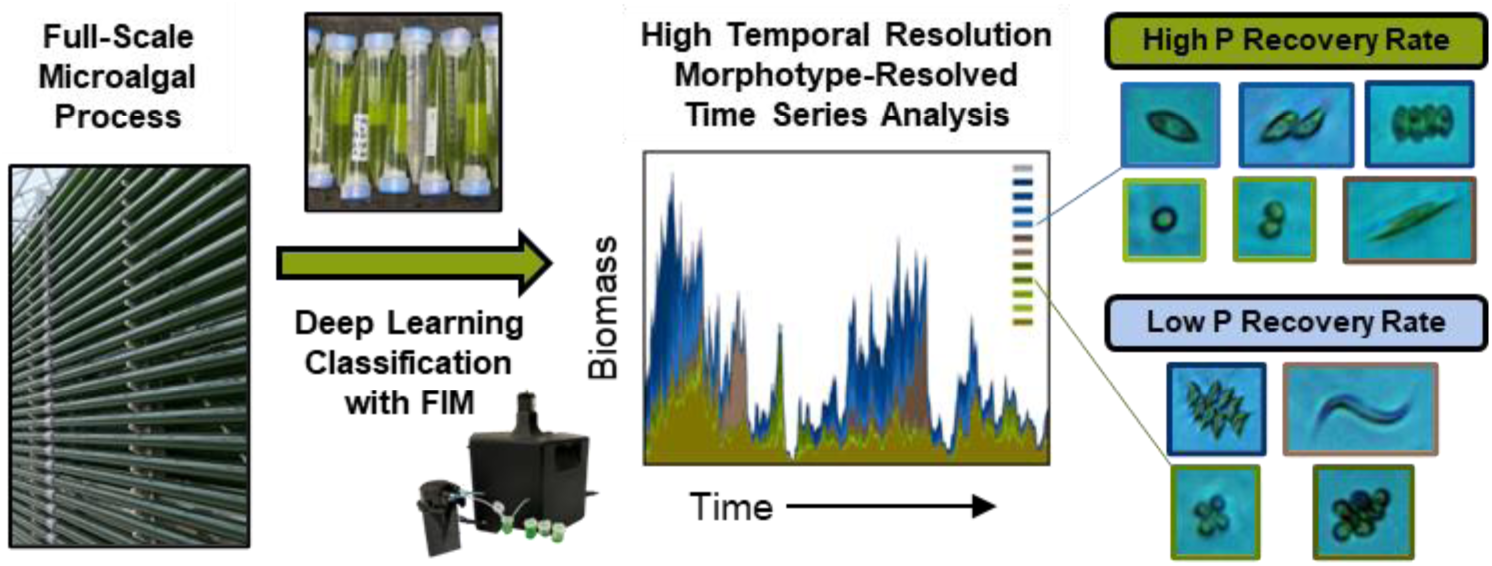

## Introduction

Increasingly strict phosphorus discharge limits across the globe (Preisner et al., 2020) are driving the adoption of microalgae-based processes for nutrient removal due to their high nutrient removal efficiency (Wollmann et al., 2019), particularly for phosphate recovery (Mao et al., 2021). Additionally, interest in microalgae’s potential to play a key role in the circular bio-economy (Corrado and Sala, 2018) has spurred the development of algae-based treatment (Lu et al., 2018) and harvesting technologies (de Morais et al., 2023). A number of microalgal cultivation formats have been adopted to meet the requirements of individual water treatment operations, ranging from large-scale lagoons (Benemann et al., 1977) to biofilm-based systems (Gross et al., 2015) to continuous cultivation in photobioreactors (Gao et al., 2016). Regardless of system design type, routine monitoring is essential to identify and mitigate disruptions to the microalgal biomass, and by extension its ability to perform the system’s designed function of nutrient recovery.

In suspension culture formats, i.e., photobioreactor cultivation systems, bulk biomass density can be monitored in real-time using optical technologies or by sampling for higher resolution analysis (e.g., gene sequencing). Total suspended solids (TSS), determined through the use of optical sensors (i.e., spectrophotometry) or manually using standardized protocols (Azema et al., 2002), is one of the key measures of biomass quantity and process health for microalgal systems. Yet, bacteria inevitably comprise a measurable portion of the system biomass even in tertiary wastewater treatment processes (Jiang et al., 2021) and TSS estimates cannot distinguish bacterial biomass from algal. While sequencing-based methods are available to resolve community composition (Alam et al., 2024), these approaches are still relatively cost-prohibitive and require a significant resource investment to allow for high-resolution data collection. Digital imaging and image processing provide a (near) real-time approach for monitoring microalgal biomass and characterizing its community composition. Various flow-through microscope systems have been developed for this purpose, in some cases with modified software tailored specifically for microalgae (Havlik et al., 2013).

Commonly utilized flow imaging microscopy (FIM) platforms for phytoplankton characterization include FlowCam (Yokogawa Fluid Imaging Technologies), Imaging Flow Cyto-Bot (McLane Research Laboratories), and ImageStreamX (Amnis EMD Millipore Sigma) (Hildebrand et al., 2016), among other commercially available instruments. Additionally, a number of low-cost alternatives have been developed in academia, including SAMSON (Deglint et al., 2018), PlanktoScope (Pollina et al., 2022), and ARTiMiS (Gincley et al., 2024), to improve access to digital microscopy for algae research. High-throughput digital microscopy platforms provide data with single-cell resolution and enable a number of downstream analyses. At their most basic, these instruments can provide a count of total particles, which can be used to infer total biomass concentration. While cell count represents a simple and unambiguous quantitative metric for population enumeration, measurement of 2D visual features (e.g., surface area) from digital microscopy has been shown to more accurately estimate total phytoplankton biomass (Mirto et al., 2015). Beyond bulk quantitation, microscopy is widely used to resolve taxonomy, enabling the enumeration of specific taxonomic groups of interest. Furthermore, the high optical resolution of these systems can distinguish discrete morphological characteristics such as colony formation in some families of green algae including *Scenedesmaceae* (Oda et al., 2022) and *Chlorellaceae* (Fisher et al., 2016; Li et al., 2018).

In this study, we aimed to evaluate the accuracy of biomass quantitation using two imaging-based platforms, FlowCam and ARTiMiS, and leveraged the unique data format that flow imaging microscope systems provided to characterize biomass composition in terms of both taxonomy and morphology. After training accurate classification models, we inferred the total abundance of taxonomic and morphological classes from a daily sampling campaign across a two-year study period. With this high resolution microalgal dataset, we explored the relationships between key taxonomic and morphological classes and the nutrient recovery performance (i.e., phosphorus removal) of a full-scale municipal wastewater treatment plant. These results demonstrated one of many high-resolution data gathering applications for characterizing microalgal systems with high-throughput imaging.

## Materials and Methods

### Sample Collection and Online Monitoring

Microalgal samples were collected daily from the Village of Roberts Wastewater Treatment Plant (Village of Roberts, WI, USA) which has been previously described (Molitor et al., 2024). Briefly, this facility utilizes microalgae for phosphorus recovery to meet a six-month average effluent total phosphorus limit of 0.04 mg-P·L^−1^. The system is comprised of several stages with internal recirculation, notably consisting of (1) a completely stirred tank reactor (mix tank) with no light availability, (2) a naturally lit tubular photobioreactor (PBR; supplemented with artificial light in low light conditions), and (3) a solid-liquid separation stage utilizing membrane filtration to decouple solids retention time (SRT) from hydraulic retention time (HRT). The facility was extensively monitored with online probes and daily or weekly on-site analysis of water chemistry analytes (pH, dissolved oxygen, temperature, orthophosphate, nitrogen species, etc.) (Molitor et al., 2024).

### Total Suspended Solids (TSS) and Nutrient Species Analysis

An online probe (ViSolid 700 IQ, YSI Inc.) recorded TSS at five-minute intervals. TSS was also manually measured on a twice-daily basis following the method described in (Bradley et al., 2021; Gardner-Dale et al., 2017). Briefly, biomass samples were filtered through 0.7 μm glass fiber filters (Whatman GF/F, Cytiva 1825142), dried at 105°C for 60 min, and desiccated for 30 min before weighing. Nutrients (NO_2_^−^, NO_3_^−^, NH_4_^+^, total nitrogen, PO_4_^3-^, total phosphorus) and alkalinity were measured in once- to twice-daily grab samples from the full-scale system using HACH colorimetric kits as described by (Molitor et al., 2024). An online orthophosphate (OP) analyzer (Alyza IQ PO_4_, YSI Inc.) recorded influent and effluent orthophosphate concentration at five-minute intervals. To correct for sensor drift and systematic erroneous readings from online probes/analyzers, manual measurements (TSS, PO_4_^3-^, etc.) were used in conjunction with probe data to construct a calibration model to reconcile errors in the probe-reported time series (Kim et al., 2024; Molitor et al., 2024).

### Flow Imaging Microscopy (FIM)

Samples for biomass characterization (total suspended solids and flow imaging microscopic analysis) were collected from a sampling port following the mix tank stage and immediately before the photobioreactor stage. Samples for on-site flow imaging microscopy were diluted between 1x to 8x (depending on current biomass concentration) for processing on a FlowCam 5000 (20X objective, 0.5 mL sample volume) flow imaging microscope operating in auto-image mode (Yokogawa Fluid Imaging Technologies, Inc.). Parallel samples were stored as 10 mL sample volume in 15 mL conical tubes, sealed with parafilm, and refrigerated at 4 °C. Daily samples were stored on-site, then shipped every two weeks with next-day shipping to Georgia Institute of Technology (Atlanta, GA, USA) for ARTiMiS imaging. ARTiMiS was configured as previously described, at 5X magnification with a 200 µm flow cell in stop-flow operation with 90-second settling times between frames (Gincley et al., 2024).

### Data Management and Accession

Water quality and system parameters (i.e., flow rate, biomass harvest, etc.) of the EcoRecover process were managed in a relational database. For variables with multiple daily measurements, values were aggregated by day (mean, range, etc.) during retrieval to align the temporal resolution of all measured quantities. FlowCam FIM feature table data was exported as .csv files per library using FlowCam’s VisualSpreadsheet software (Version 5.7.19), and corresponding particle images were extracted from the onboard relational database directly through an unofficially supported program. ARTiMiS feature table data and images were directly retrieved from the instrument’s file system. ARTiMiS sample metadata, feature table data, and images were managed in an external graph database, and particle data were aggregated by class label during retrieval for daily quantity measurements.

### Total System Biomass Analysis

Total biomass estimates from online TSS probe, ARTiMiS, and FlowCam were correlated with manual TSS estimates; manual TSS estimates were considered the benchmark due to the implementation of a recognized standard protocol (Lipps et al., 2023). Given the five-minute temporal frequency of TSS probe data, daily mean was used to match the temporal frequency of other modalities for comparison. FIM data from FlowCam and ARTiMiS were normalized on a per-sample basis by aggregating various metrics of interest (e.g., total count, particle area, etc.) and scaling by the sample dilution factor divided by the total sample volume to yield a metric-per-unit-volume quantity (e.g., total surface area per μL). Each modality was compared on a day-by-day basis, and days lacking one or more modalities for a particular date were discarded before regression. Outliers were identified based on interquartile range (IQR), with discard thresholds defined as (3 x IQR) < Q_1_ and (3 x IQR) > Q_3_ where Q_1_ and Q_3_ refer to the 25^th^ and 75^th^ percentiles, respectively. Linear regression was used to estimate the degree of fit between each measurement pair, with manually measured TSS as the predicted variable for fitting. ARTiMiS FIM data were binned by focus label (in- or out-of-focus), assigned by a pre-trained binary neural network classifier, to determine the effect of including only in-focus particles as compared to including all detected particles to estimate biomass concentration.

### Class Library Annotation

Annotation libraries corresponding to classes of interest (the most abundant taxonomic groups and their common morphology types where applicable) were initialized by collecting representative examples of each class until a minimum critical quantity (approx. 100 individual particles) was annotated. Further annotation of each class was then expedited using each FIM technology’s respective computer-assisted sorting functionality. For FlowCam: new sample runs were loaded, and particles were filtered by feature match score to the target library. Individual particles with high match scores were manually annotated into the respective class library. For ARTiMiS, a prototype convolutional neural network (CNN) classifier was trained to identify the target class(es) and applied to unseen sample data to predict the class identity of individual particles, which were then automatically sorted into class-labeled directories. These prediction folders were manually curated by reassigning labels for misclassifications before merging with the parent annotation library. This process was repeated iteratively (model training, inference, and misclassification refinement) until a minimum of 750 and up to 2000 unique particles were included in each class library. Priority was given to classes yielding low test accuracy scores during manual refinement. As an artifact of this computer-assisted annotation process, feature distributions of FlowCam libraries were observed to be normally distributed within discrete decision boundaries due to the explicit statistics-based sorting in the VisualSpreadsheet software, whereas ARTiMiS libraries were not.

### Particle Classification Models

FlowCam’s native analysis program, VisualSpreadsheet, includes built-in classification functionality. However, this system operates using a set of thresholds on individual semantic features (e.g., diameter, sphericity, etc.) applied in sequential combinations for each class in a classification template; this produces a high misclassification rate for different classes with overlapping feature distributions. Given the current wide availability of robust machine learning and deep learning classification techniques, these default VisualSpreadsheet methods were avoided. Instead, annotated library data were extracted from the FlowCam database for use in external classification techniques for analysis. Two deep learning classification model types were used: a multi-layer perceptron (MLP) with two hidden layers utilizing feature table data as input (hereby referred to as a deep neural network (DNN)), and a convolutional neural network (CNN) utilizing region of interest (ROI) images as input, both providing class predictions as output. Model architectures used are slightly modified versions of the “micro neural network” models described in (Gincley et al., 2024). Models trained on ARTiMiS-derived data were of identical architecture to, and trained from scratch using the same training protocol (number of epochs, cross validation, etc.) as, the FlowCam data-based models. To serve as a reference, a MobileNetV2 classification model (Sandler et al., 2019) was trained using transfer learning on the same dataset; upstream blocks were frozen and the final prediction output block was modified to match the number of classes tested here.

### Classification Model Training Methodology

For image-based classifiers (CNNs), regions of interest (ROIs) were grayscaled then down-sampled or padded to a uniform shape of 128×128×1 pixels. Each unique ROI was augmented through a complete set of mirrors and rotations to yield 8x images per ROI for training; approximately 12,500 unique ROIs were transformed into a training dataset of approximately 100,000 ROIs. For feature table-based classifiers (i.e., DNNs), approximately 50 features were used for training. Feature table data was not augmented to avoid generative data transformations requiring assumptions about feature distributions. As a result, the sample size depth to input variable width ratios were approximately 6.1 for CNNs and 250 for DNNs, respectively. Training data depth and breadth were consistent for both FlowCam and ARTiMiS training datasets. All datasets were holdout-split to designate distinct training and test sets before augmentation and cross-validation, with a test split ratio of 15%. Following augmentation, training data were split into training and validation sets during three iterations of Monte Carlo cross-validation (random resampling each iteration). For CNN classification models, training data were divided into six batches and trained for 10 epochs in each batch (60 epochs per cross-validation iteration, 180 epochs in total). Models were trained on an NVIDIA 3060 Ti graphics processing unit (GPU) with 8 GB video memory and 32 GB system memory. The DNN classification models were trained without dataset-batching for 15 epochs each CV (45 epochs in total). Epoch values for each model type were selected based on plateaued performance on the validation set in preliminary testing and fixed as described for consistency across experiments. Models were iteratively retrained from scratch to yield a “best” classifier based on a composite score maximizing overall accuracy and minimizing inter-class accuracy variability.

### Time Series Shifted Correlation Analysis

All analytes included in correlation analysis were cleaned of outliers via the IQR-based method as described earlier and then normalized by scaling [0,1]. Sparsity in individual time series was filled via linear interpolation with daily resolution, and local extrema were smoothed with a three-day rolling mean (the minimum window size for a centered moving average). Datasets were truncated to predetermined start and end dates for specific time windows of analyses (e.g., high and poor nutrient removal performance periods). To infer predictability (i.e., statistical causality) from results, individual pairwise analytes were procedurally time-shifted up to a specified maximum time lag before calculating Spearman’s rank order correlation coefficients. This approach was taken under the mechanistic assumption that interactions in the system were not instantaneous, but rather were governed by a temporal lag. Given the assumption that the exact duration of this lag would vary on an analyte-pair basis, precedent for an elastic algorithm was taken from the field of financial modeling (Stübinger, 2019; Stübinger and Adler, 2020). Significant relationships were defined as follows: for time lags between three and ten days, if a majority (4) of the correlations met the significance criteria (p < 0.05), the relationship between the correlated parameters was considered significant. The correlation statistic representing each correlated pair was selected as the maximum observed among the significant individual lags. Three days was selected as the minimum lag time under the design requirement of predictive relationships being actionable during routine operation, i.e., proactive measures could be taken to rectify trajectories.

## Results

### Biomass estimations with online probes and FIM

To evaluate the accuracy of individual modalities to estimate total biomass, manually measured TSS (mTSS) was selected as the ground-truth measurement. Online TSS as estimated by a ViSolid 700 IQ TSS probe (pTSS) was compared against FIM-based measurements to determine whether the additional level of detail provided by FIM improved biomass estimation. Samples used for this analysis spanned two years of data collection and ranged from 0-800 mg·L^−1^ (mTSS). Raw pTSS readings were corrected (see *Materials and Methods*) to compensate for sensor drift and other sources of error before comparative analysis. Linear regression was selected as a modeling technique to capture how effectively each parameter modality served as a proxy measurement for mTSS.

A simple linear regression between corrected pTSS measurements and mTSS values resulted in a RMSE value of 110 mg·L^−1^ with a coefficient of determination R^2^=0.62, representing a “best-case” scenario for the use of probe measurements (**Figure 1A**). This served as a reference for FIM-based biomass estimation which is the focus of this investigation. Two broad methodologies were available to estimate total system biomass from FIM-based particle data: count-based measurements and area-based measurements. Count-based measurements refer to the number of objects (i.e., cells and colonies) detected. In contrast, area-based measurements refer to the surface area (size) of particles as measured in pixels. Linear regression of individual FIM-based measurements to estimate mTSS indicated that area-based measurements more accurately predicted mTSS than count-based measurements for both FIM technologies (**Figure S1**). The best (lowest RMSE) measurement type for each FIM technology is shown in **Figure 1**. The “total filled particle area” metric was determined to be the lowest-error predictor measurement from FlowCam data (**Figure 1B**), which represents the summed foreground-mask pixel areas of all particles for each respective sample. Simple linear regression to mTSS yielded an RMSE of 142.4 mg·L^−1^ with coefficient R^2^=0.43. The lowest-error measurement type from ARTiMiS data was similarly determined to be the summed total particle surface area metric (**Figure 1C**), again representing the total pixel areas of all particles observed in each sample. Simple linear regression to mTSS yielded an RMSE of 98.3 mg·L^−1^, the lowest error rate observed in this study, with coefficient R^2^=0.75, the highest measured. The ARTiMiS analytical pipeline automatically labeled individual objects as either in- or out-of-focus. Thus, it was possible to determine to what extent particle focus had impact on biomass estimation. It was found (**Figure S1**) that aggregation of only out-of-focus particles resulted in the highest biomass estimation error rates among ARTiMiS measurements. Interestingly, this estimation error was still lower than that of the lowest-error (i.e., best) FlowCam-based measurement type. Aggregation of both in- and out-of-focus labeled objects resulted in lower error than using only in-focus particles, which suggested that enumerating out-of-focus objects provided informational value for biomass estimation even in cases where these particles might not be identifiable. These results also decisively established that total particle area provided a better estimate for total biomass than particle counts regardless of imaging system.

**Figure 1:**
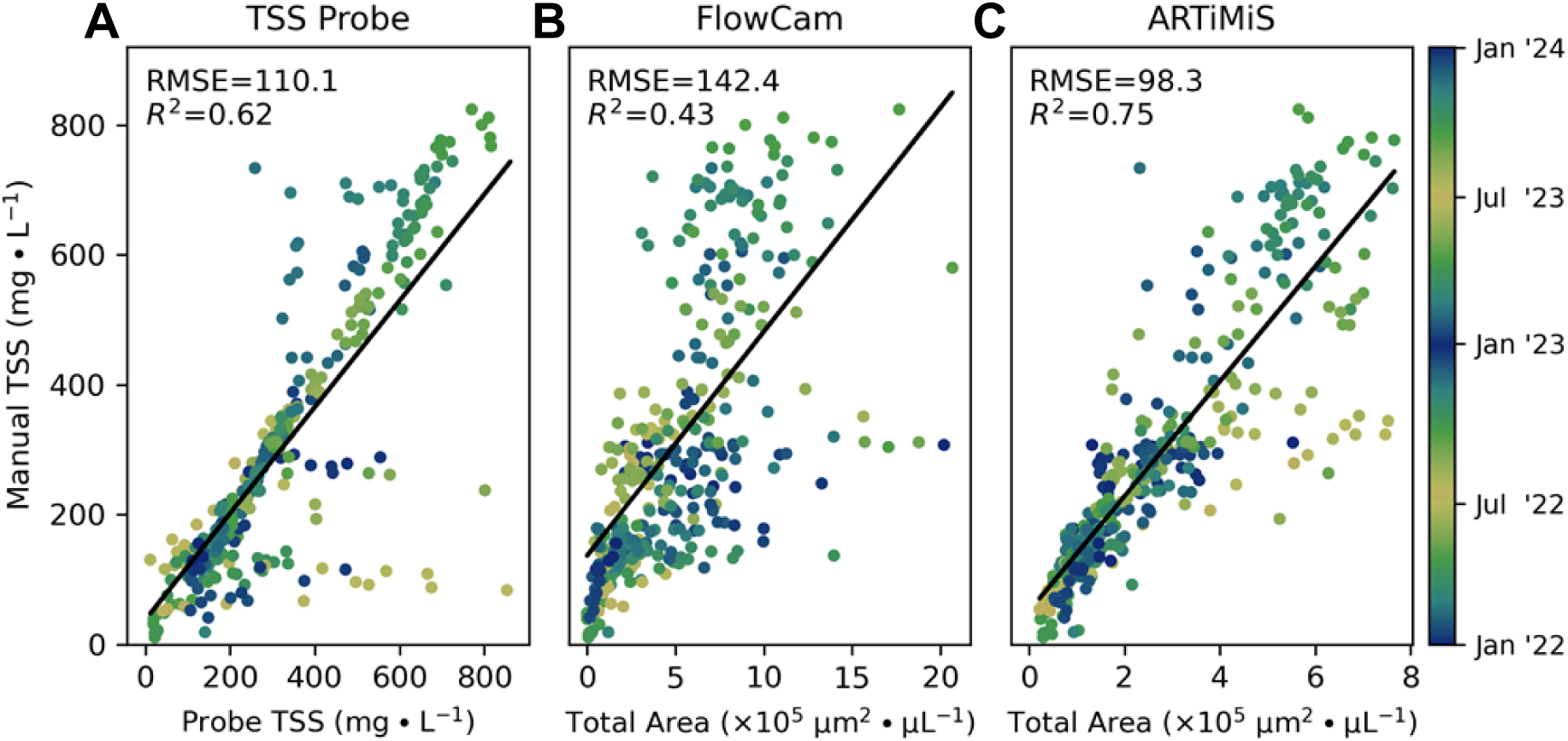
Comparison of total biomass estimation methods benchmarked to manual TSS. These methods include **(A)** calibrated probe TSS data, and total particle surface area as a proxy for biovolume as measured by **(B)** FlowCam and **(C)** ARTiMiS instruments. Individual samples are colored temporally to represent seasonal trends in total system biomass.

### Classification from semantic particle features

From raw image data and refined particle feature data, objects were identified as described previously (Gincley et al., 2024; Mirasbekov et al., 2021). Feature data in tabular format represented the lowest-dimensionality information provided by both FlowCam and ARTiMiS, and thus theoretically enabled the use of the most lightweight classification models (fewest trainable parameters). In addition to taxonomic identity, variation in cell morphological type (i.e., morphotype) for each taxon were represented by independent classes.

Libraries of particles belonging to the dominant taxonomic groups were annotated, subdivided by specific morphological types and/or species labels where applicable, and were used to train deep neural networks (DNNs) of identical architecture to predict class identity based on their respective feature table data. Thus, this analysis aimed to determine the relative quality of particle measurements and annotation libraries derived from the two instruments. Feature distributions of morphotype subclasses of the same taxonomic group were characteristically different between libraries generated from the two instruments (**Figure 2A, 2B**). For example, the 4-cell and 8-cell variants of *Scenedesmus* exhibited minimal overlap with other variants in the FlowCam data. More extensive feature overlap was observed in ARTiMiS libraries, which has the potential to impact closed-set accuracy during model training and testing. To train the classification models, annotation libraries were subdivided into training data and test data subsets, and each classifier was evaluated on the unseen test data subset to estimate its generalized performance. It is important to note that although the unique particles in the test dataset did not overlap with any training data, these data were nonetheless subject to the same, if any, annotation biases present in the complete dataset. When evaluated on the unseen test dataset, the FlowCam data-based classifier (**Figure 2C**) notably outperformed the ARTiMiS data-based classifier (**Figure 2D**). This result remained consistent when morphotypes were aggregated at the taxon level (**Figure S2**). These results suggested that the FlowCam-derived particle features more robustly represented their distinct classes when compared to ARTiMiS-derived particle features; for a given DNN classifier complexity, training dataset size, and training intensity, they produced classification models with higher test accuracy. However, these results were suspected to have been influenced by the class separation observed in the annotated dataset features, and thus may not be representative of accuracy in un-annotated contexts.

**Figure 2:**
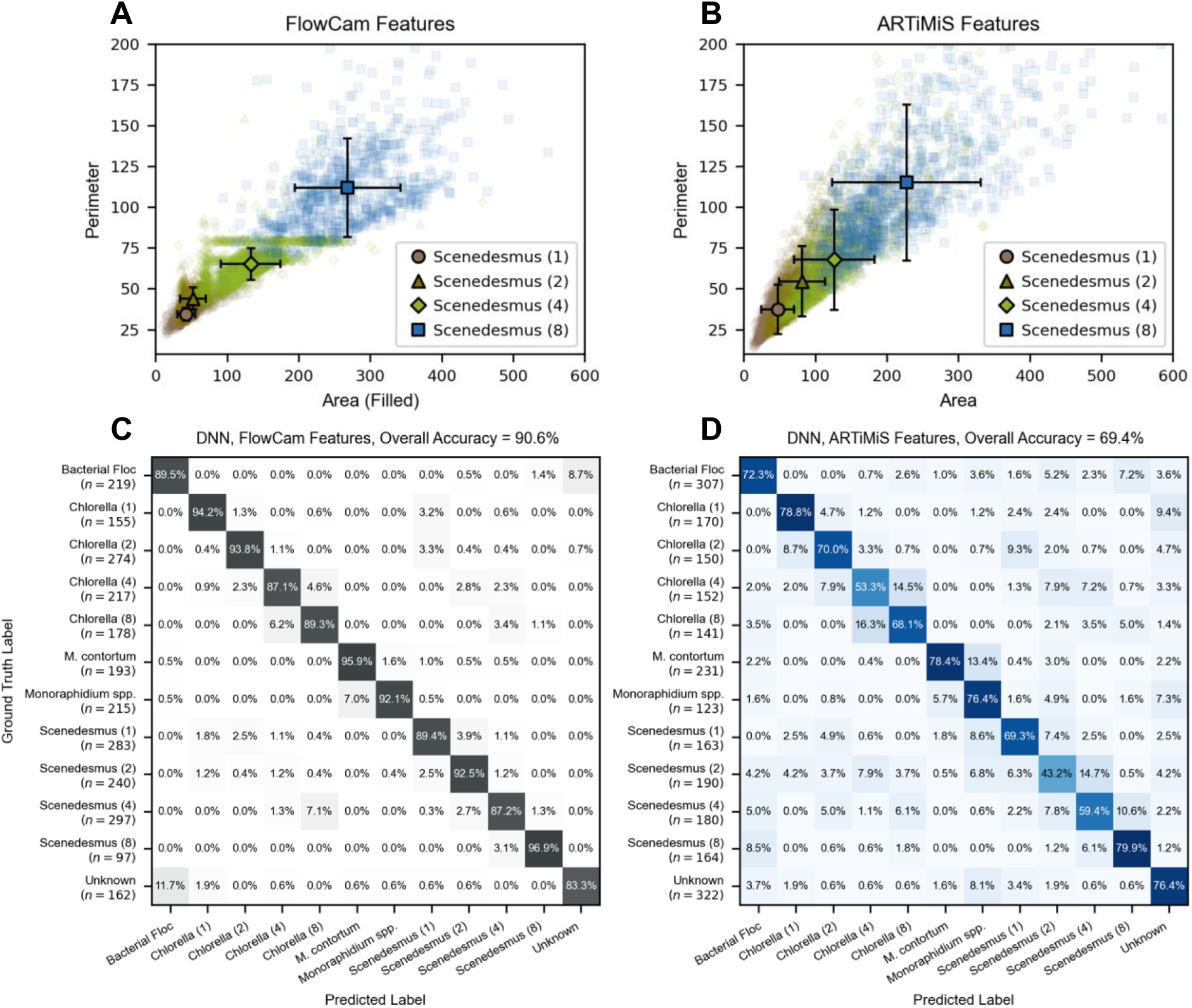
Distribution patterns of two important semantic features for particle recognition from **(A)** FlowCam-based annotated image libraries and **(B)** ARTiMiS-based libraries. Classes are distinct morphological types of microalgae in the *Scenedesmus* genus. Individual particles are shown as scatter points, population mean, and standard deviation are shown as bolded points and bracketed bars. Performance of DNN classification models trained on a subset of the particles depicted in **(A, B)** and tested on an unseen subset of the same libraries, depicted as confusion matrices. Number of test samples indicated as *n* for each class, representing 15% of the total annotated library size used for training. On-diagonal values indicate correct prediction frequency, matrix coordinates indicate pairwise misclassification rates. DNNs of the same architecture correctly predict taxonomic and morphotype identity with 90.6% overall accuracy when trained on FlowCam Data **(C)**, and 69.4% accuracy when trained on ARTiMiS data **(D)**.

### Classification from direct particle images

Previous work (Gincley et al., 2024) evaluating classification of microalgae from the EcoRecover system suggested that image-based classification with convolutional neural network (CNN) classifiers outperformed feature table-based classifiers (i.e., random forest, DNNs). Utilizing revised CNN model architecture and model training routines, and expanded annotation libraries, CNN classification performance was re-assessed in this study. This same classification approach was implemented with FlowCam-captured image data retrieved from the same class libraries as **Figure 2**. In doing so, the objective was to provide a comparison of image data quality for use in classification between the two FIM platforms.

Classification with a CNN using image data improved both per-class and overall classifier accuracy for both FlowCam-derived (**Figure 3C**, **Figure S3A**) and ARTiMiS-derived (**Figure 3D**, **Figure S3B**) test datasets for both morphotype-resolved and taxonomic class-collapsed labels. As with DNN-based classification (**Figure 2**), FlowCam-based data resulted in higher classification accuracy compared to ARTiMiS-based data, though this gap was significantly narrower. These results were obtained using a micro-CNN architecture (Gincley et al., 2024); to determine if the publicly available and significantly larger MobileNetV2 CNN classifier could recover the accuracy gap, the same ARTiMiS dataset was used to train this model using transfer learning. While taxon class-collapsed classifier accuracy was comparable to the micro-CNN (**Figure S3D)**, MobileNetV2 underperformed on the morphotype classification task (**Figure S3C)**.

**Figure 3:**
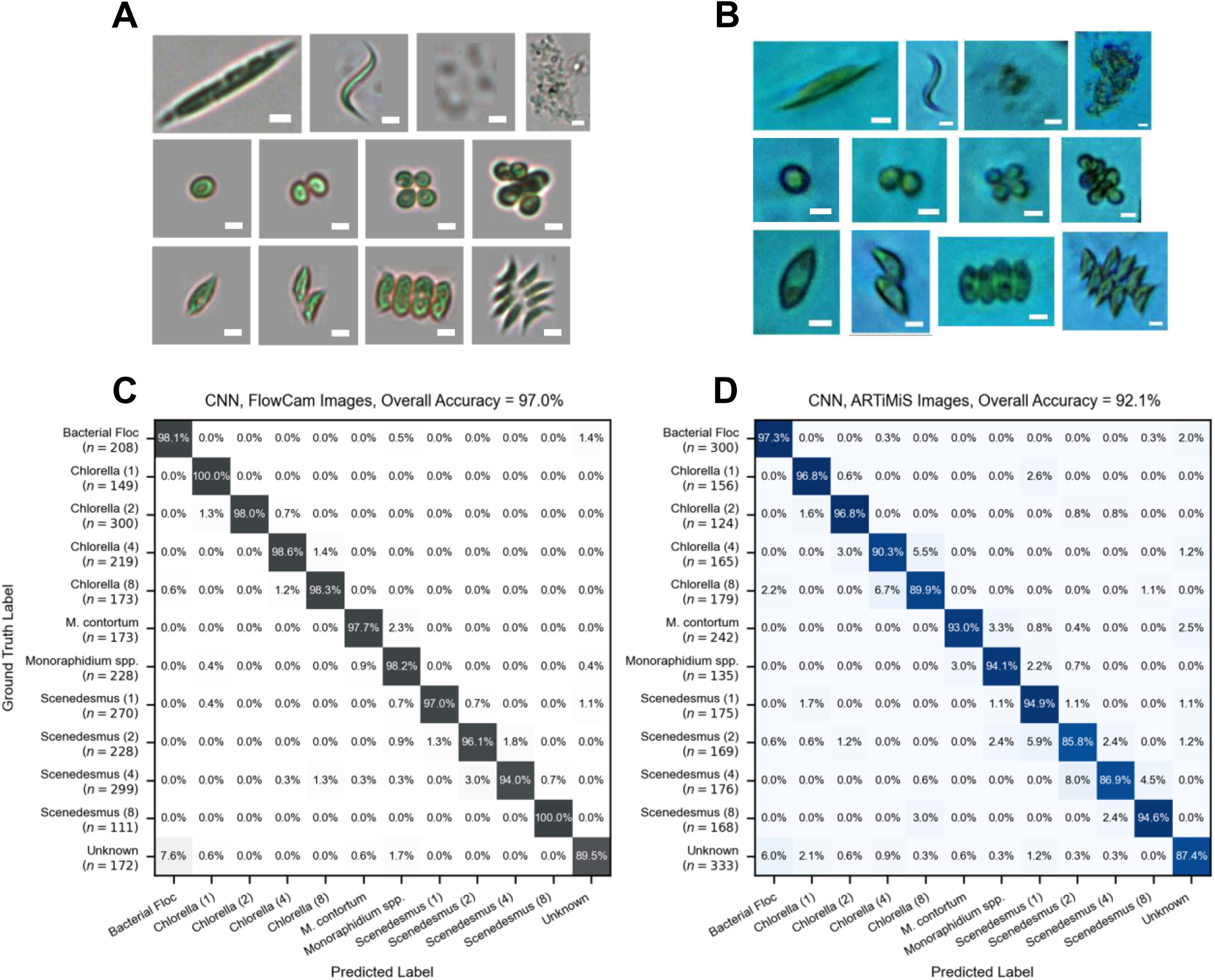
Sample images of morphologically distinct variants of dominant taxonomic groups captured on FlowCam **(A)** and ARTiMiS **(B)**. Top Row: *Monoraphidium spp.*, *Monoraphidium contortum*, Unknown particle class, Bacterial Floc class. Middle Row: *Chlorella spp.* spanning 1-, 2-, 4-, and 8-cell morphological variants Bottom Row: *Scenedesmus spp.* spanning 1-, 2-, 4-, and 8-cell morphological variants. Scale bars = 5 μm. Confusion matrices from CNNs of the same architecture trained to distinguish between microalgae classes from libraries of annotated images captured on FlowCam **(C)** and ARTiMiS **(D)**. Sample size of each class in test dataset indicated by *n*.

### Observational analysis of time series data

After training accurate classification models, the best-performing ARTiMiS-based CNN classifier was used to estimate the abundances of the classes of interest across the complete study period, from January 2022 to December 2023 (two complete years). This resulted in the generation of a time series dataset with single cell resolution, each assigned a predicted class label. Populations could be viewed as a total event count, or any other attribute aggregate: namely, total surface area per class per sample point. These aggregations could further be represented either as absolute or relative magnitude (**Figure S4**), each providing a unique view of biomass composition. Since the absolute total surface area was determined to be the most accurate representation of system biomass (**Figure 1**), this was selected as the primary view of microalgae populations over time (**Figure 4A**). A time series of microalgae population dynamics with high temporal resolution enabled direct comparison with the EcoRecover system’s key process performance indicator (i.e., effluent orthophosphate (PO_4_^3-^)) to identify relationships between microalgal populations and system performance. Effluent orthophosphate (OP) was characterized both as a percentage of removal (**Figure 4B**) and as absolute concentration (**Figure 4C**), using measured influent OP as a reference. Qualitative inspection of the time series suggested that total biomass quantity was related to OP removal efficiency (rapid declines were often aligned with low removal rates), but also indicated that the composition of such biomass may have had an impact on OP removal as well. For example, during the period of May-June 2022, OP removal rate degraded significantly despite relatively high total system biomass; one key biological difference was the dominance of *Monoraphidium* during this period, displacing the *Scenedesmus* population typically observed in the microalgal community. From this time series, several high and poor performance periods were identified for closer inspection, being operationally defined as periods consistently meeting phosphorus discharge permit requirements (e.g., December 2022 to February 2023) and consistently exceeding permit requirements (e.g., June to August 2022), respectively.

**Figure 4:**
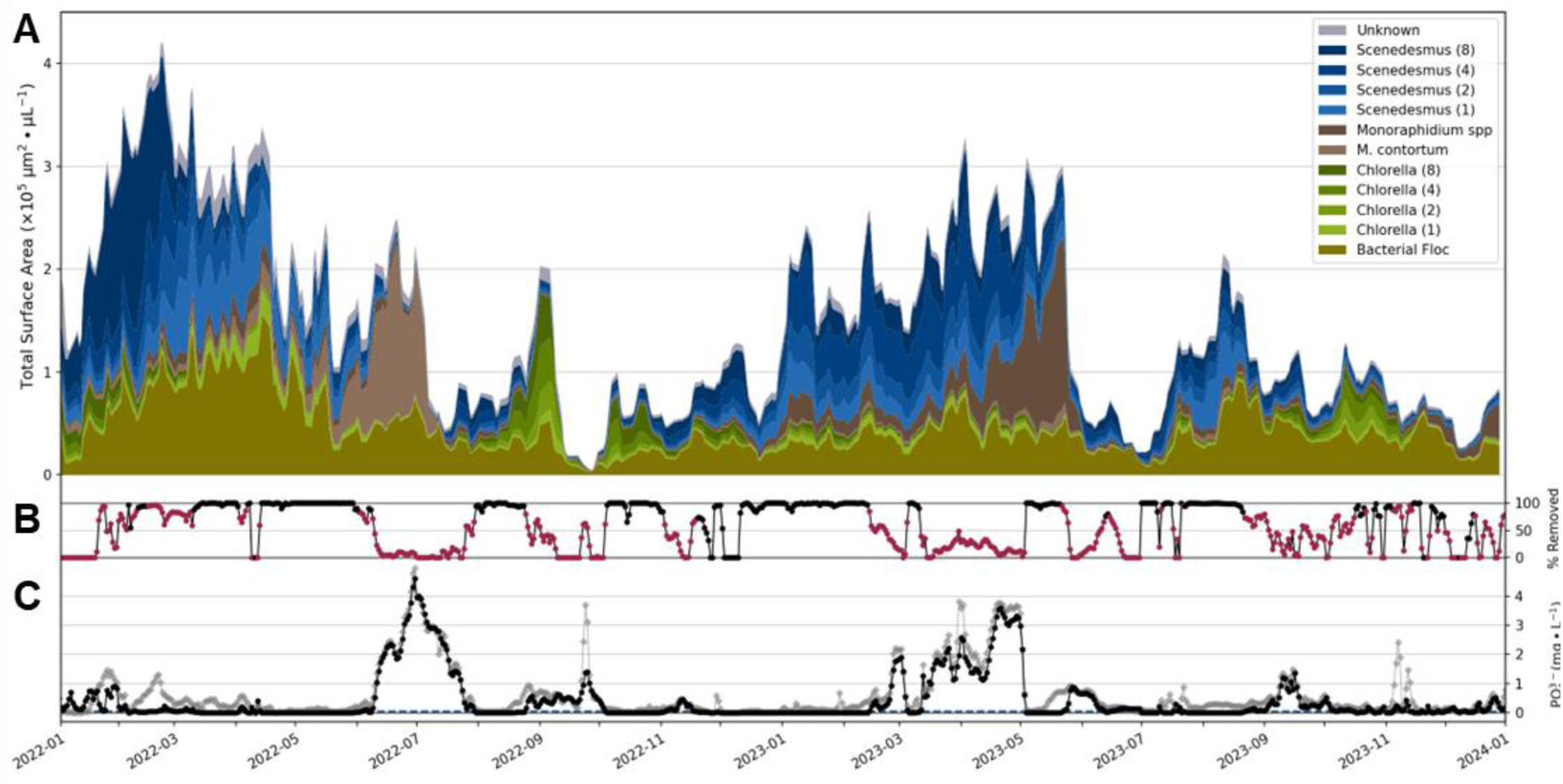
**(A)** Time series depiction of microalgal biomass represented as total area per unit volume. Daily measurements of each particle class stacked to sum to total biomass. **(B)** OP removal with samples exceeding permit level indicated in red. **(C)** Time series of total influent (gray diamonds) and effluent (black circles) orthophosphate (OP) measured, and corresponding percent OP removed. Local permit level of 0.04 mg-P·L^−1^ indicated by dashed navy blue line.

### Correlation analysis of time series data

The qualitative observation that community composition appeared directly related to system performance prompted a statistical investigation to identify significant relationships between the microalgae and key process parameters. A hybrid statistical approach was implemented to infer time-dependent correlations between specific analyte pairs. Specifically, a time-lag method was applied while utilizing Spearman’s rank order correlation coefficient as the test statistic. Individual analyte pairs were time-lagged between three and ten days and assessed for statistical significance. Lagged correlations were assessed during specific temporal periods associated with poor orthophosphate removal (**Figures 5** and **S5**), high orthophosphate removal (**Figures 6** and **S6**), and for the duration of the complete study period (**Figure S7**). The period of poor orthophosphate removal spanning May-July 2022 (**Figure 5**) was attributed largely to microalgal community composition as the cause of degraded performance, in combination with extremely high influent orthophosphate loading (peaking at over 4 mg·L^−1^) during this period. The total system biomass was observed to be generally stable during this period despite a consistently deteriorating orthophosphate removal rate culminating in a major process failure in late-June 2022 (**Figure 4**). The increase in the concentration of *Monoraphidium contortum* was accompanied by a decrease in the concentration of all other microalgae populations. This displacement of *Chlorella* and *Scenedesmus* likely drove the reduction in phosphorus uptake, nearing 0% removal. This change in community composition coincided with increases in (1) temperature, (2) influent nitrate concentrations, and (3) influent orthophosphate concentration, and decreases in (1) influent ammonium concentrations and (2) influent alkalinity levels. Periods of poor performance were not always directly related to microalgal taxonomic succession, as observed from May to June 2023 (**Figure S5**). Immediately prior to this period, the population of non-specific *Monoraphidium spp*. increased, however OP removal did not appear to be impacted until a sudden and extreme decline in total microalgal biomass occurred. This sudden event was attributed to (1) high temperatures, (2) mild acidification, and (3) depletion of total nitrogen in the EcoRecover system which exhibited a strong negative correlation with lagging microalgae populations. The rapidly degraded system performance was therefore not due to a succession of the *Monoraphidium* population in this case, but rather its collapse.

**Figure 5:**
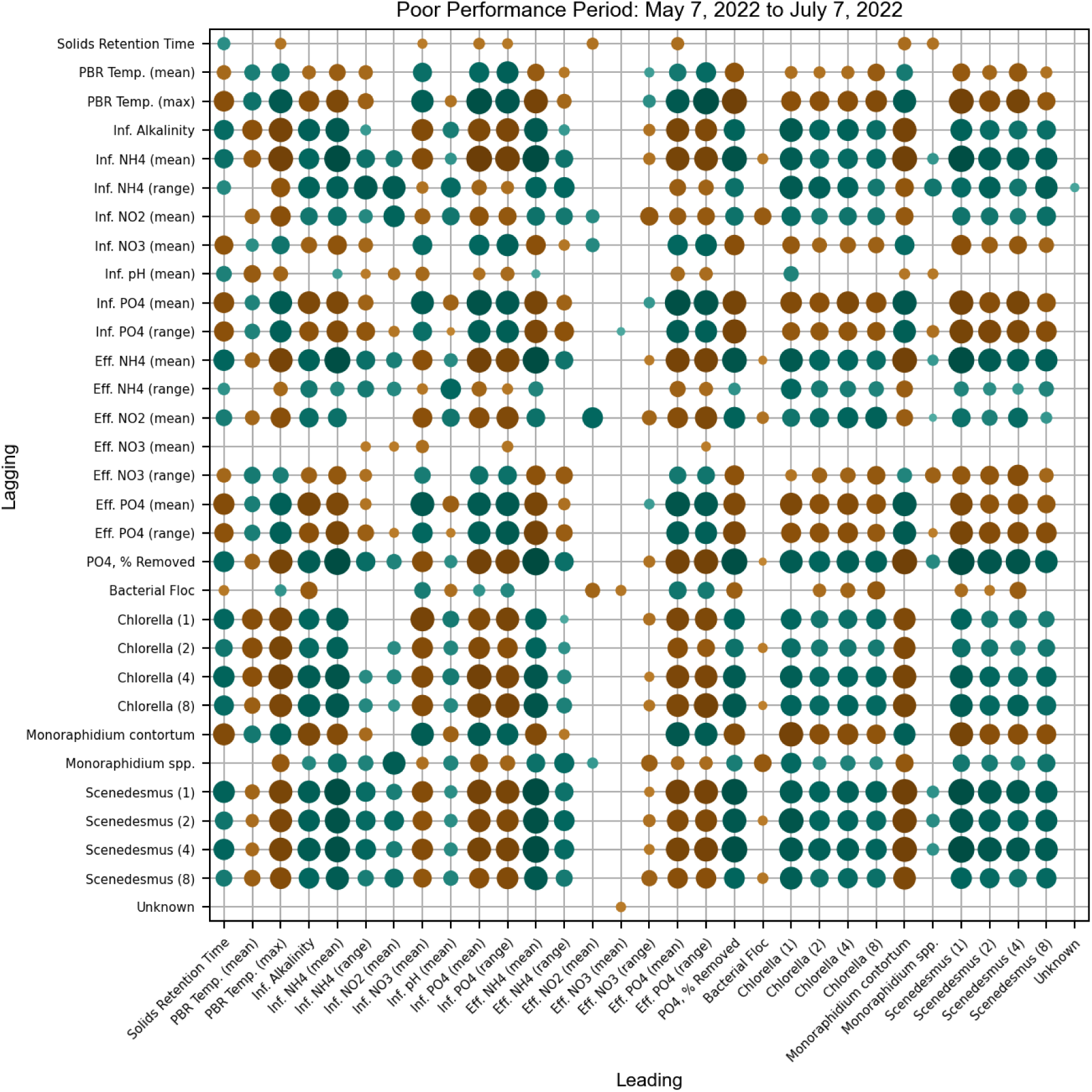
Asymmetric correlation matrix of key EcoRecover system parameters, including microalgae biomass as total area, during a period of poor performance (low orthophosphate recovery rate). In pairwise analysis, variables on the vertical axis have a time-lag with the variables on horizontal axis. Mean and range quantities describe statistics on a per-sample-day basis (i.e., daily range). Influent abbreviated as Inf.; Effluent as Eff. Spearman rank correlation coefficients are indicated by size and color: green = positive correlation, brown = negative correlation. Positive correlation describes time series that move in the same direction (e.g., both decreasing), negative correlation describes movement in opposite directions (e.g., one increasing, other decreasing). Indices without statistical significance (p < 0.05) omitted.

**Figure 6:**
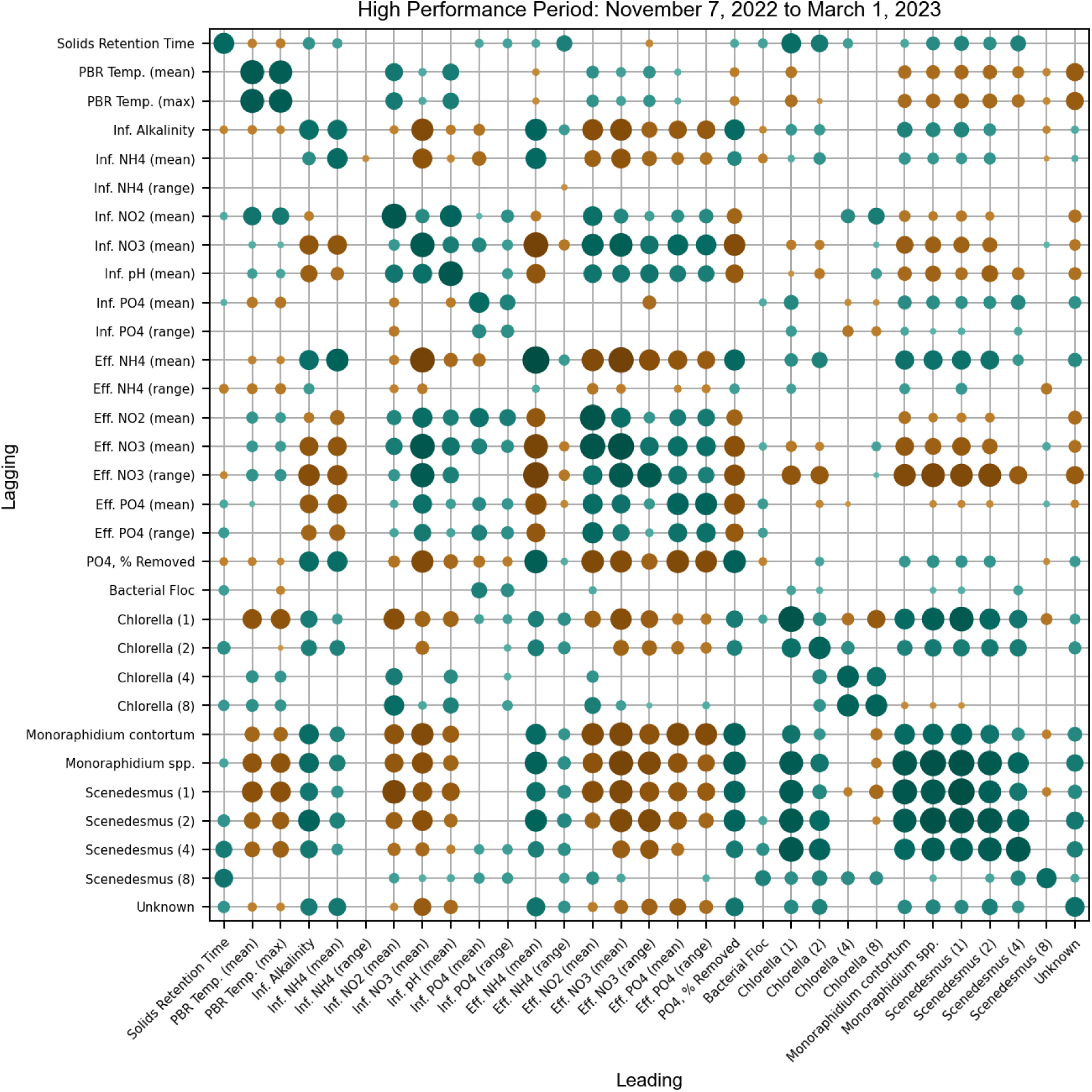
Asymmetric correlation matrix of key EcoRecover system parameters, including microalgae biomass as total area, during a period of high performance (high orthophosphate recovery rate). In pairwise analysis, variables on the vertical axis have a time-lag with the variables on horizontal axis. Mean and range quantities describe statistics on a per-sample-day basis (i.e., daily range). Influent abbreviated as Inf.; Effluent as Eff. Spearman rank correlation coefficients are indicated by size and color: green = positive correlation, brown = negative correlation. Positive correlation describes time series that move in the same direction (e.g., both decreasing), negative correlation describes movement in opposite directions (e.g., one increasing, other decreasing). Indices without statistical significance (p < 0.05) omitted.

Investigation of high system performance periods (**Figure 6** and **S6**) indicated that OP removal was driven largely by microalgae in the *Scenedesmus* genus, specifically the smaller (1- and 2-cell) morphotypes, and the *Monoraphidium* genus. Smaller (1- and 2-cell) morphotypes of *Chlorella* were also weakly positively associated with OP removal. Interestingly, larger morphotypes of *Chlorella* were negatively associated with OP removal and appeared to be driven by increasing temperatures and decreasing influent pH. The microalgal taxa positively associated with OP removal were observed to be positively impacted by increased ammonium concentrations and to a lesser degree by longer solids retention times. In an effort to determine more universal relationships between system analytes, the complete study period was assessed (**Figure S7**). Overall, the microalgae population appeared to be negatively impacted by increasing temperatures and nitrate concentrations, and positively impacted by increasing alkalinity and ammonium concentrations. As expected, OP removal rate was weakly positively associated with all green algae studied, with *Scenedesmus* most consistently appearing as important for high system performance. While most microalgal populations auto-correlated with one another, another notable outcome of this analysis was the mild negative association between large-morphotype *Chlorella* and the *Monoraphidium* classes studied, potentially implying either a competitive relationship or a divergence in optimal growth conditions.

## Discussion

### ARTiMiS-based total biomass estimates were most consistent with manual measurements

ARTiMiS FIM yielded biomass estimates that reflected mTSS values with lower error rates and higher linearity than both pTSS and FlowCam FIM estimates, suggesting that it may serve as a more effective technology to automate routine TSS monitoring. ARTiMiS estimates were significantly more consistent (lower residual variance) than FlowCam, which was unexpected given both systems utilize the same fundamental data collection method. Furthermore, inspection of the reported values of total surface area per unit volume from the two FIM platforms (**Figure 1B, 1C**) revealed that FlowCam systematically over-reported surface area relative to ARTiMiS. One potential explanation for these differences could be attributed to the rate of background residual particle accumulation during FlowCam sample runs. At the start of each sample run the instrument initialized a calibration image to identify objects adhered to the flow cell, sometimes referred to as artefacts, i.e., background noise to ignore. However, it was observed that this automated system was either unable to account for new background objects accumulated during the sample run, or its background object selection criteria were too strict to account for variability in position or appearance resulting from interactions with the sample matrix during the course of a sample run (or both); not a unique challenge among FlowCam instruments (Álvarez et al., 2012; Camoying and Yñiguez, 2016; Marvin et al., 2019).

Due to the high volume of narrow field of view (FOV) individual frames captured (typically ∼7,000 per sample in this study), thousands of “false positives” – physically-duplicate but not feature-identical objects – were recorded as unique ROIs, impacting aggregate sample statistics. Importantly, the lack of feature identity (e.g., the same object observed at different coordinates in sequential frames) posed barriers to algorithmic de-duplication. Though these objects were able to be manually deleted in post-processing to some extent, this technique was inherently challenging to perform accurately, consistently, and completely. The amount of total background noise was therefore impacted by sample composition, instrument maintenance state, and manual artefact removal, which added uneven variability over the two-year study. While ARTiMiS did not perform a background particle removal step in this study, a combination of flow cell material properties (resulting in a low particle adhesion), fluid handling protocols (high velocity flushing between frame captures), and a high-FOV, low-frame count operational design were implicated in contributing to a significantly lower incidence of background residual particles affecting sample results. For low-biomass applications where background noise has a more pronounced impact, a tailored adaptive solution has been reported (Khan et al., 2024).

Upon further examination of the mTSS approximations, ARTiMiS-based measurements were normally distributed about a mean of zero regardless of total biomass, whereas pTSS and FlowCam-based data exhibited non-normal or non-zero mean residual distributions (**Figure 1**). This observation was quantitatively supported by the comparatively low RMSE values for ARTiMiS measurements. Importantly, this suggested that ARTiMiS provided more consistent predictions across a wide range of biomass concentrations with normally distributed errors, indicating that it could estimate mTSS biomass with more predictability in a variety of conditions (i.e., proportional margins of error at 60 mg·L^−1^ and 600 mg·L^−1^). For these reasons, the more consistent ARTiMiS data was selected for biomass characterization in time series analysis.

### Particle surface area was shown to more accurately model biomass than particle counts

Both FIM technologies recorded an array of measurements on a per-particle basis, including calculations for predicted biovolume based on geometrical assumptions (particle is spherical, spheroid, etc.). These derived quantities were calculated from surface area and related morphological features (e.g., major and minor axis length). In order to reduce errors resulting from these assumptions about 3D geometry (Mirto et al., 2015; Owen et al., 2022), the primary measurements (surface area) were used directly for biomass estimation. It is also important to note that the influent to the EcoRecover system was secondary wastewater effluent, containing bacteria. Total system biomass, captured in measurements of TSS, was comprised of this microalgal-bacterial consortium (Jiang et al., 2021). The use of particle surface area, rather than cell count, therefore provided a more accurate picture of the composition of microalgal and bacterial biomass fractions (**Figures 4**, **S4**), and represented another distinct advantage of using FIM to accurately characterize biomass compared to more traditional solids-based methods.

### Particle classification using deep learning provided accurate identification

FIM instruments like FlowCam and ARTiMiS generate both raw images and refined morphological features (e.g., particle area, circularity, etc.) as exportable data. From these datasets, machine learning models have the potential to automate routine tasks including classification. Such classification techniques have, by now, been widely explored in phytoplankton research (Ciranni et al., 2024), and are becoming established as standard procedure to make use of the large quantities of data generated by FIM platforms. In this study, we compared the relative accuracy of feature table-based and image-based deep learning classification models using data from both FIM platforms. A DNN classifier trained on FlowCam feature table data outperformed the same type of classifier trained on ARTiMiS data (**Figure 2**). This result was likely due to several factors but may be primarily attributable to the structure of the annotated class library feature distributions (**Figure 2A, 2B**). This was likely caused by differences in how computer-assisted annotation was performed on each platform (see *Materials and Methods*). A side-effect of the FlowCam annotation method was that population outliers were less likely to be included during the annotation workflow to increase library sizes, as these particles were excluded by the filtering process. Distinct separations between classes can be observed at approximately one standard deviation from the population means (**Figure 2A**). In contrast, the borders between ARTiMiS-based library classes (**Figure 2B**) appear less structured. This annotation bias observed in the test set suggests that the FlowCam classifier is more likely to be less accurate when applied to true un-annotated raw sample data (which will invariably contain a noteworthy incidence of outliers and edge cases), whereas the ARTiMiS result may be interpreted as a more realistic approximation for real-world performance.

### CNN-based classification demonstrates equivalent accuracy from ARTiMiS and FlowCam image data

CNN classification models have been shown to provide improved accuracy over feature table-based classifiers, particularly for visually complex tasks (i.e., microalgae discrimination) (Gincley et al., 2024). This was found to be true in this study as well, with CNN classifiers for both FIM platforms out-performing the DNN models. In the case of CNN classification, ARTiMiS performed nearly as well as FlowCam, and given the previously identified side-effects of annotation bias in the datasets, the gap between the two systems is likely narrower in real-world applications. This gap in accuracy is narrowed further when class labels are binned by taxonomic group (**Figures S3A, S3B**), where ARTiMiS was shown to achieve 95.7% overall accuracy in a taxonomy classification task. These results are significant given ARTiMiS’ order-of-magnitude lower instrument cost, demonstrating that capital-intensive FIM platforms are not required to achieve high identification accuracy.

The CNN classifier architecture used to achieve these results (for both FIM platforms) was designed to reduce computational costs by minimizing total trainable model parameters (200K parameters in this study). This was done to decrease the number of training samples and total training time required, as well as to optimize compatibility with low-power hardware (Gincley et al., 2024). This second point was critical to enable real-time classification onboard the ARTiMiS instrument which runs on a single-board computer. This “micro-CNN” architecture was benchmarked against MobileNetV2 (Sandler et al., 2019), a larger (2M parameters) publicly available CNN designed for edge computing applications. The “micro-CNN” was seen to out-perform MobileNetV2 for this classification task (**Figure S3**). This is important to note, given the propensity for contemporary researchers to rely on large “off-the-shelf” CNN models to perform complex phytoplankton classification tasks (Eerola et al., 2024; Lin et al., 2022; Yang et al., 2023), and suggested that improved performance (both in terms of accuracy and computation time) may be achieved with smaller, task-tailored CNN architectures.

### High-resolution temporal monitoring illuminates factors affecting process performance

The study period consisted of a dynamic, diverse collection of system states: periods of complete orthophosphate removal, and of incomplete removal; periods of high and low biomass; periods of a stable, stratified microbial community and periods of complete takeover by singular taxonomic groups. Several of these periods were isolated for analysis (**Figures 5**, **S5**, **6**, **S6**), and the full time series was processed to elucidate any generalized trends (**Figure S7**). The behavior of the dominant chlorophytes in the EcoRecover system (*Chlorella*, *Monoraphidium*, and *Scenedesmus*) was of particular focus across these specific periods and across the full study period, and were examined individually to compare observations in this system to other contemporary microalgae-based wastewater treatment systems. In the EcoRecover process, *Scenedesmus* appeared to be one of the key taxa associated with high system performance. *Scenedesmus* has been consistently reported (Morillas-España et al., 2021; Mubashar et al., 2024; Silambarasan et al., 2023) to efficiently remove phosphorus from wastewater, which coincides with observations from this study. It has recently been reported that a *Chlorella* isolate was the dominant taxonomic group for nutrient removal in WWTP effluent (Baldisserotto et al., 2023), with further evidence of *Chlorella*’s potential for P assimilation from synthetic wastewater (Asaad and Amer, 2024). In contrast, *Chlorella* in the EcoRecover system appeared to have limited P-removal efficacy based EcoRecover system performance data (**Figures 4, S5, S6**).

While it has been reported that cold-tolerant *Monoraphidium* can be used to treat wastewater effluent (Kirchner et al., 2022), the *Monoraphidium* taxa observed in this study appeared to coincide with warmer overall temperatures, whereas *Scenedesmus* was the dominant group during colder months. *Monoraphidium contortum* has previously been examined as the dominant chlorophyte for wastewater treatment in one study (Choi et al., 2023), in which nitrogen supplementation was necessary for optimal biomass growth. Importantly, a higher biomass was recorded in that study with a low N:P ratio of 1.7:1. This finding may partially explain the succession observed in early summer 2022 as *M. contortum* grew to over 90% of the chlorophyte biomass, as the EcoRecover influent N:P ratio dropped precipitously mid-June from over 100:1 to under 10:1 and remained low until early August. Previous literature (KV et al., 2018) indicates that *M. contortum* can sustain unperturbed growth rates with limited nitrogen availability as long as phosphorus is not growth-limiting, which may explain why this taxon displaced the *Scenedesmus* population.

Few studies have examined the morphological composition of these taxa, making comparison to contemporary literature challenging. Nonetheless, the data in this study suggested that the dominance of smaller morphotypes was more positively associated with high system performance, and the abundance of larger morphotypes may be interpretable as unfavorable environmental conditions for efficient nutrient uptake for both *Chlorella* and *Scenedesmus* taxa. Emerging consensus in literature suggests that formation of larger colonies is associated with physiological stress (Albini et al., 2019; Dong et al., 2018; Fisher et al., 2016; Lürling and Van Donk, 2000). These results are consistent in that the EcoRecover system performed more poorly when its microalgae exhibited stress indicators.

The dataset collected during this two-year study was constituted of measurements with high accuracy and high temporal resolution. Despite focused analysis of both specific temporal periods and the complete study duration, definitive universal markers of expected high and poor performance remained elusive. The EcoRecover system exhibited a high degree of dynamic behavior, not uncommon in other full-scale microalgal wastewater treatment systems. Due to their highly dynamic nature, nuanced details of biological and chemical processes in these systems may only be observable with high temporal resolution. Different periods of both high performance and poor performance were driven by different factors upon close inspection, indicating that generalized conclusions from lower-resolution datasets may be misleading. High-frequency sampling, made automated by online sensors and digital FIM platforms, might help provide a more accurate picture of similar systems in future studies. Specifically, attending not only to taxonomy but also morphology is suggested to be critical to understanding the role and behavior of microalgae in such systems. Discrete morphological variation, i.e., colonial behavior, in chlorophytes is often specifically attributed to pest and/or predator pressure in contemporary literature (Albini et al., 2019; Dong et al., 2018; Fisher et al., 2016; Lürling and Van Donk, 2000). Conversely, it may also emerge from multiple fission cell division, which is described to occur during optimal growth conditions (Bišová and Zachleder, 2014), i.e., no predator/pest pressure. While this work focused on the microalgal constituency of the EcoRecover biomass, FIM also enables the detection of pest and predator organisms at relatively low abundances (Wang et al., 2017). By leveraging high-throughput, high-resolution morphotype characterization, further investigation of the interactions between microalgae and pests/predators may reveal new insights into these dynamics and could help explain some of the results of this and related studies.

## Conclusions

In this study, FIM was shown to accurately serve as a (near) real-time proxy measurement for total biomass, outperforming estimates provided by a total suspended solids probe. ARTiMiS FIM provided more accurate TSS estimates than FlowCam. Utilizing curated image databases from both FIM platforms, CNN-based classification models predicted both the taxonomic and morphological identity of microalgal and bacterial particles with high accuracy. This enabled the creation of a high temporal resolution time series dataset with both high taxonomic and morphotypic resolution. Examining this microalgal time series data alongside high-resolution water chemistry and system performance measurements highlighted significant relationships. Specifically, (1) *Scenedesmus* spp. appeared to be most important for high system performance (i.e., P removal), and (2) elevated temperatures and nitrite/nitrate concentrations appeared to have the greatest negative impact on microalgal biomass, and thus on system performance. These results suggest that high-resolution biomass characterization, made possible by FIM platforms such as ARTiMiS, can reveal key insights into system performance and process control in industrial scale microalgal processes.

## Acknowledgements

The authors would like to acknowledge the staff operating the Village of Roberts EcoRecover process for their on-site support enabling this sampling-intensive study: Village of Roberts Director of Public Works, John Bond, and the entire Public Works staff, as well as the team at Clearas Water Recovery including Kevin McGraw, Jordan Lind, and Patrick Kelly. This work was funded by the U.S. Department of Energy, Office of Energy Efficiency and Renewable Energy, under Award Number DE-EE0009270. This work was also supported by the U.S. National Science Foundation Graduate Research Fellowship Program under Award Number 2039655 and the Paul L Busch Award for Innovation in Applied Water Quality Research. Any opinions, findings, and conclusions or recommendations expressed in this publication are those of the author(s) and do not necessarily reflect the views of the U.S. Department of Energy.

