## Supplementary Information for "Morphotype-Resolved Characterization of Microalgal Communities in a Nutrient Recovery Process with ARTiMiS Flow Imaging Microscopy"

***Supplementary Figure S1:***

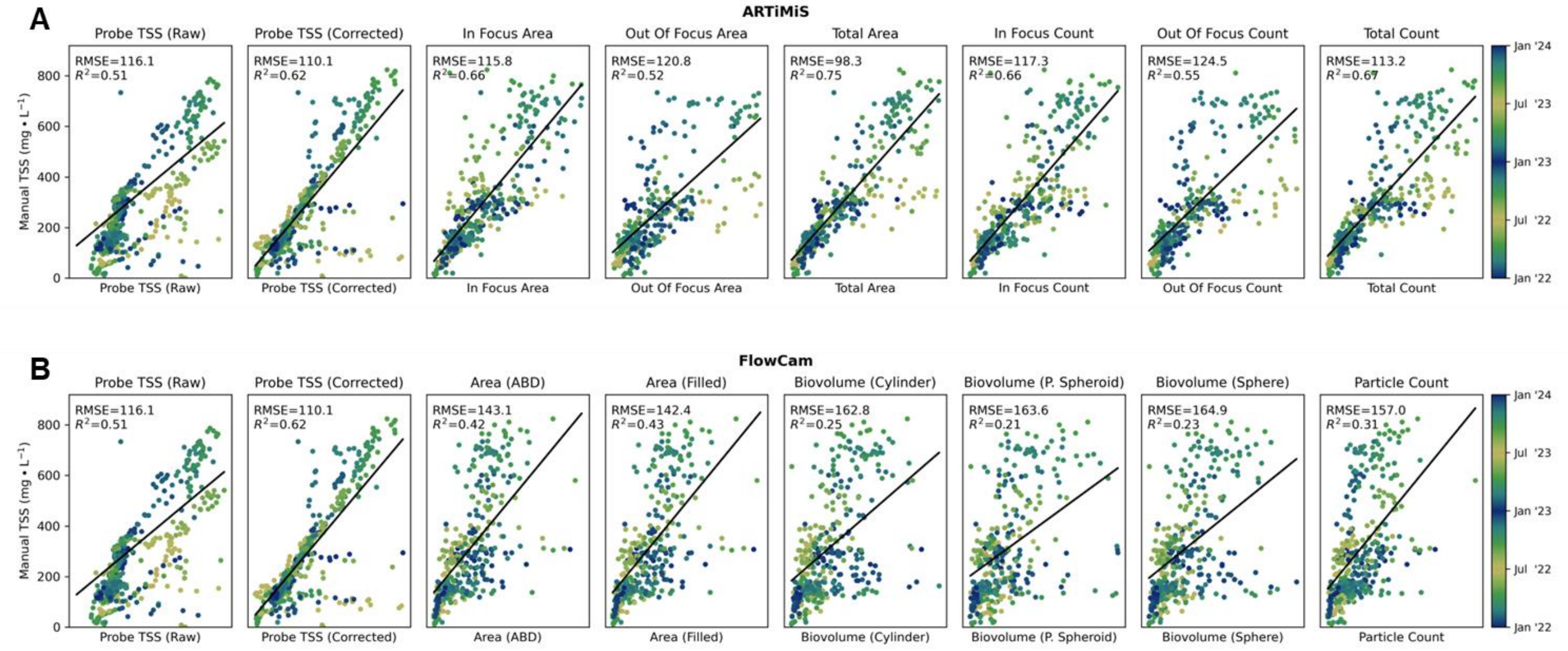


**Figure S1:** Approximations of manual Total Suspended Solids (mTSS) measurements by digital instruments. Comparison of raw and corrected TSS Probe measurements against (A) ARTiMiS FIM measurements and (B) FlowCam FIM measurements. (A) ARTiMiS automatically labels particles as in- or out-of-focus; in-, out-of-focus, and both (total) populations were enumerated both in terms of total particle surface area and total particle counts. (B) FlowCam provides multiple measurements of area: calculated from area-based diameter (ABD) or as the filled foreground pixel area. FlowCam also provides 3D biovolume estimations using assumptions of particle geometry (cylindrical, spheroid, and sphere-shaped objects).

***Supplementary Figure S2:***

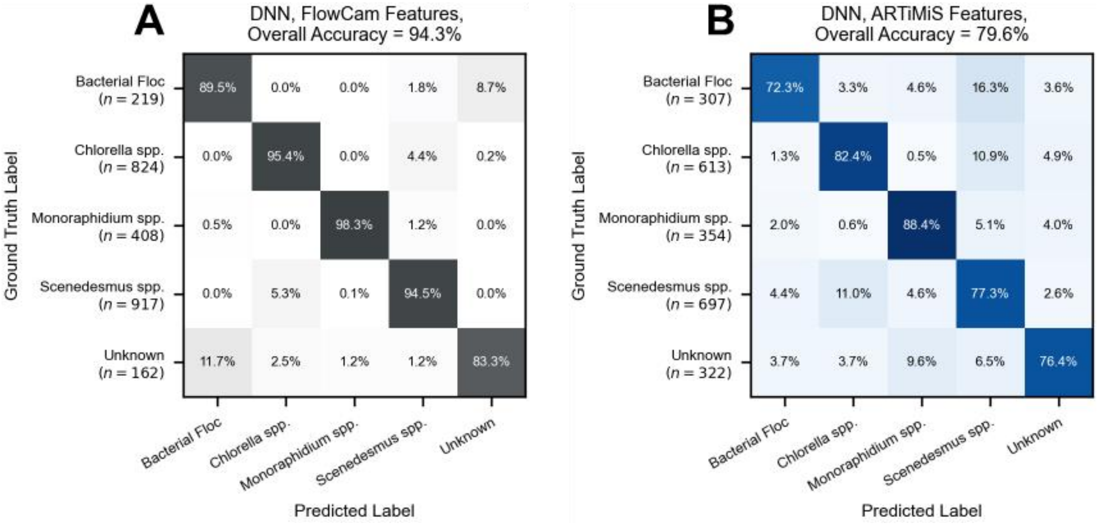


**Figure S2:** Taxonomy-collapsed deep neural network (DNN) classification model predictions from **Figure 2**, trained on feature table data. Morphotype/species labels were binned by taxonomic group and correct/incorrect predictions were scored. DNN trained on FlowCam features (A) is the same model as in **Figure 2A**. DNN trained on ARTiMiS features (B) is the same model as in **Figure 2B**.

***Supplementary Figure S3:***

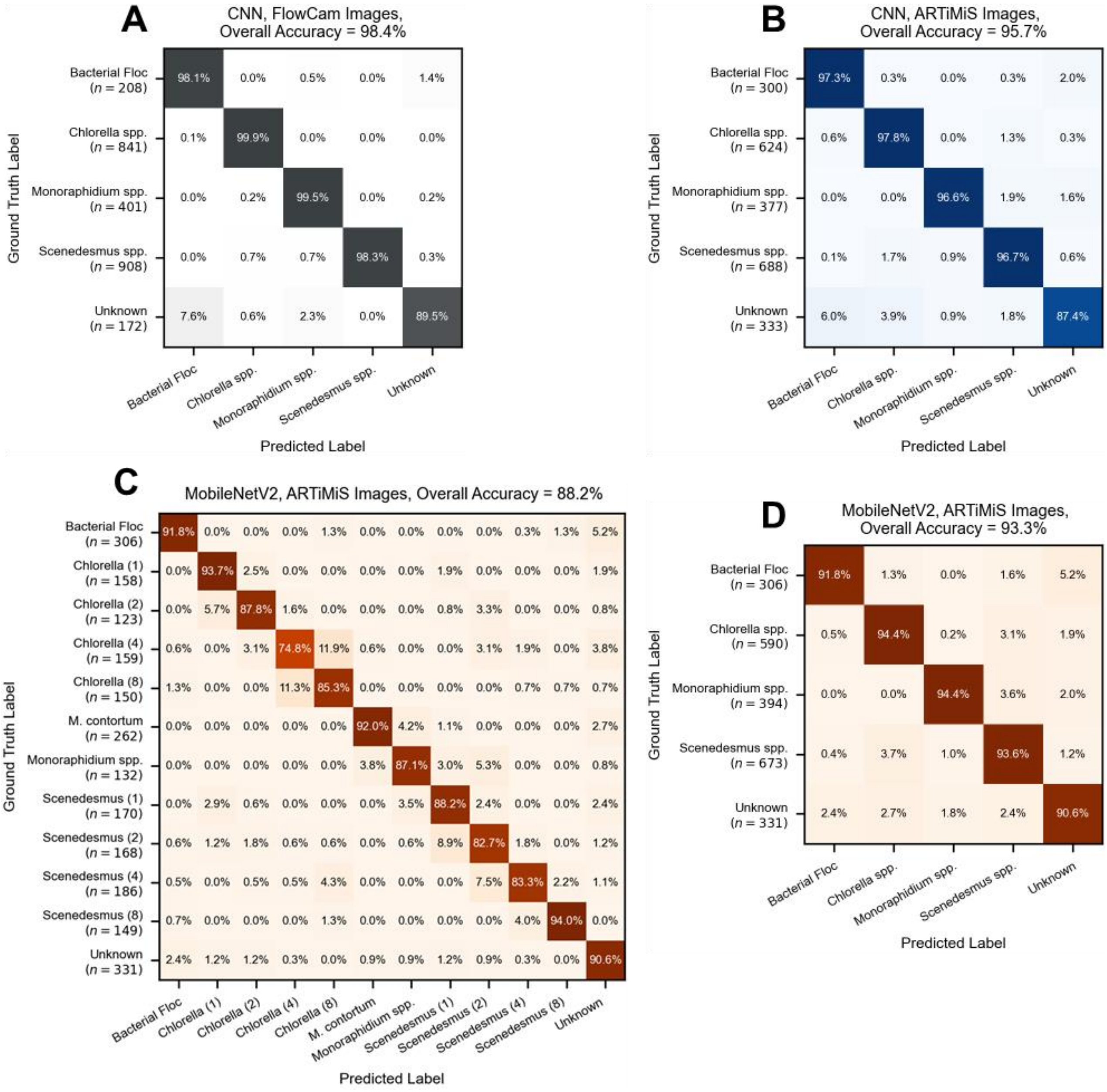


**Figure S3:** Taxonomy-collapsed convolutional neural network (CNN) classification model predictions from **Figure 3**, trained on image data. Morphotype/species labels were binned by taxonomic group and correct/incorrect predictions were scored. CNN trained on FlowCam features (A) is the same model as in **Figure 3A**. DNN trained on ARTiMiS features (B) is the same model as in **Figure 3B**. An off-the-shelf MobileNetV2 model was trained using transfer learning on the same dataset and at the same training intensity (number of epochs and cross-validation) as the ARTiMiS CNN and evaluated on the same test dataset. The morphotype-resolved (C) and taxonomy-collapsed (D) results are compared to **Figure 3** and **Figure S3**, respectively.

***Supplementary Figure S4:***

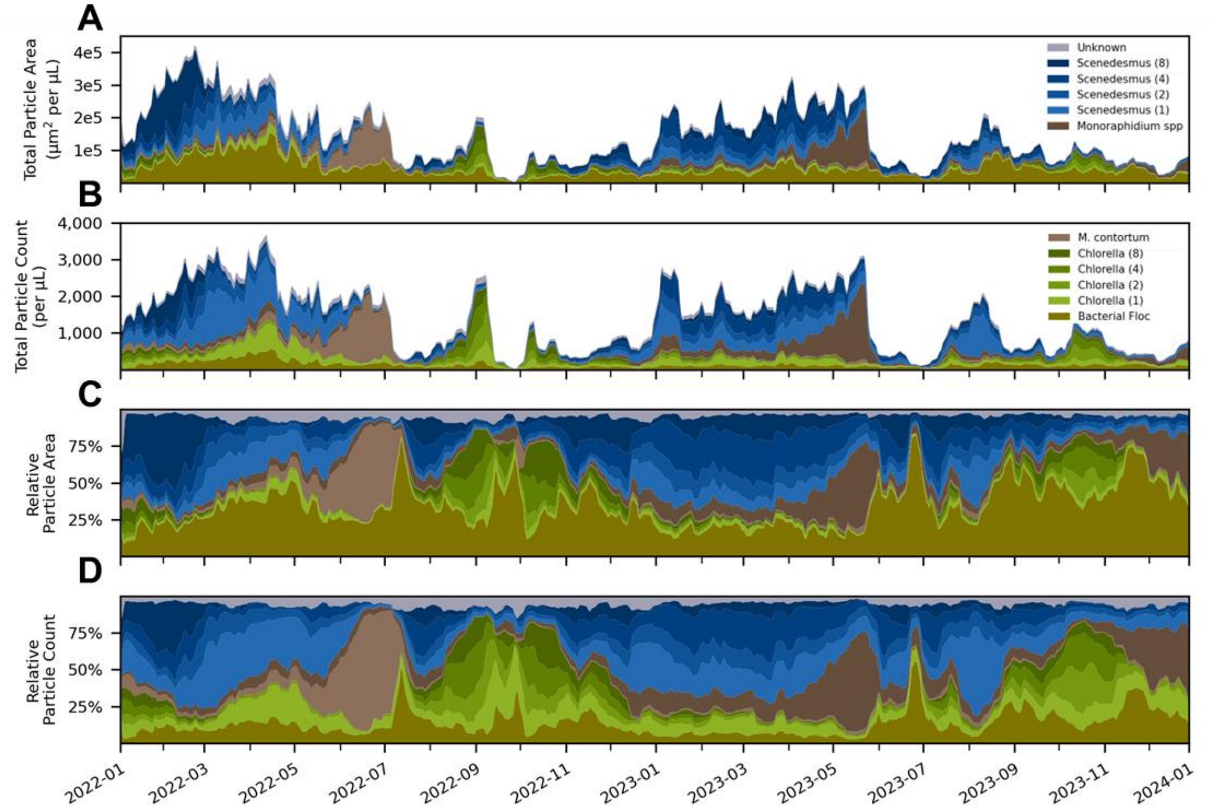


**Figure S4:** Time series views of microbial biomass depicting the four combinations of total (upper panels) and relative (lower panels) particle surface area (above) and particle count (below) view types. Colors are identical to **Figure 4**.

*Supplementary Figure S5:*

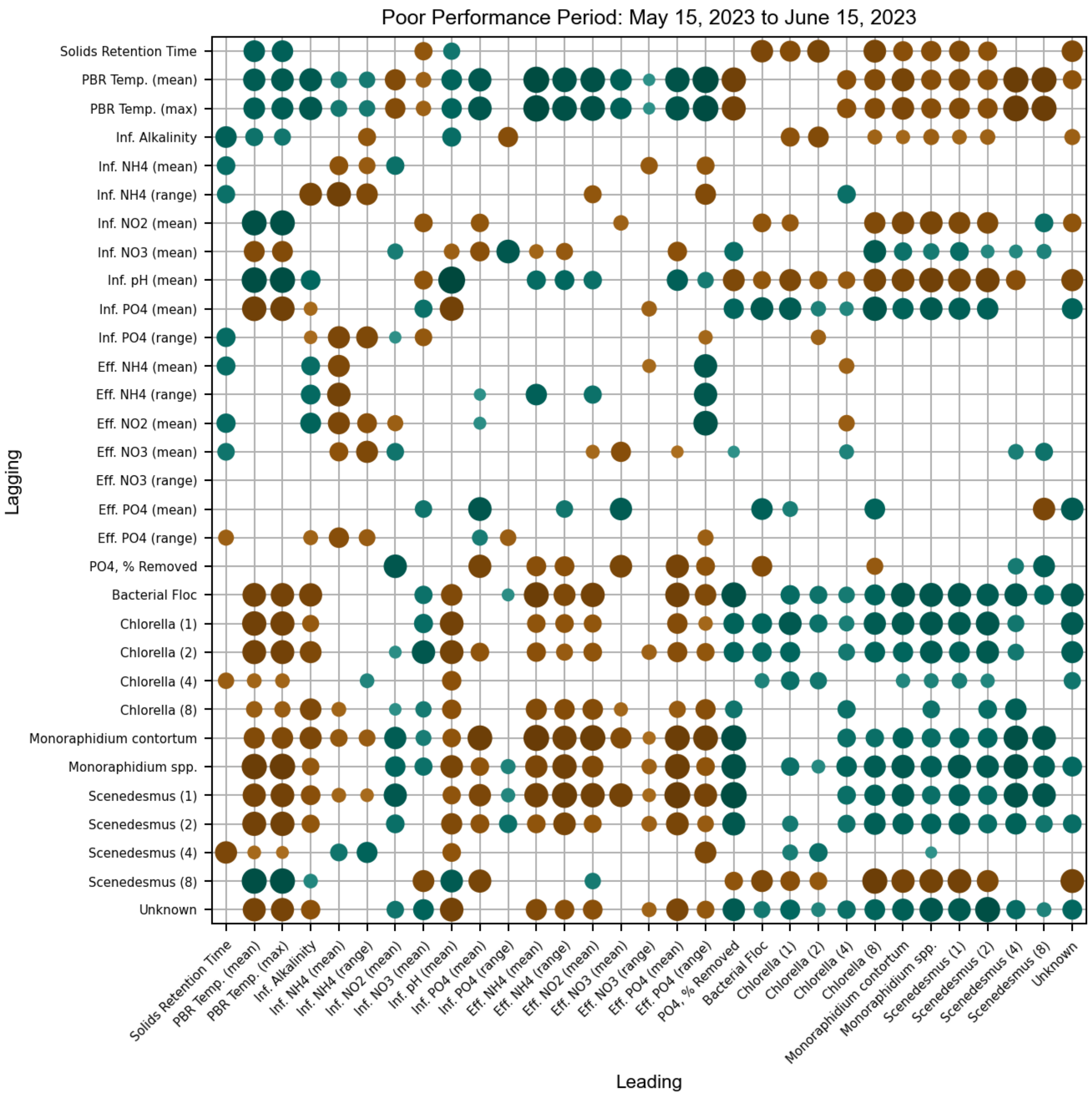


**Figure S5:** Asymmetric correlation matrix of key EcoRecover system parameters during a period of poor performance (low orthophosphate recovery rate). In pairwise analysis, variables on the vertical axis have a time-lag with the variables on the horizontal axis. Mean and range quantities describe statistics on a per-sample-day basis (i.e., daily range). The view of microalgae biomass (**Figure S4**) selected for analysis is of total area. Spearman rank correlation coefficients are indicated by size and color: green = positive correlation, brown = negative correlation. Positive correlation describes time series that move in the same direction (e.g., both decreasing), and negative correlation describes movement in opposite directions (e.g., one increasing, other decreasing). Indices without statistical significance ($p < 0.05$) are omitted.

***Supplementary Figure S6:***

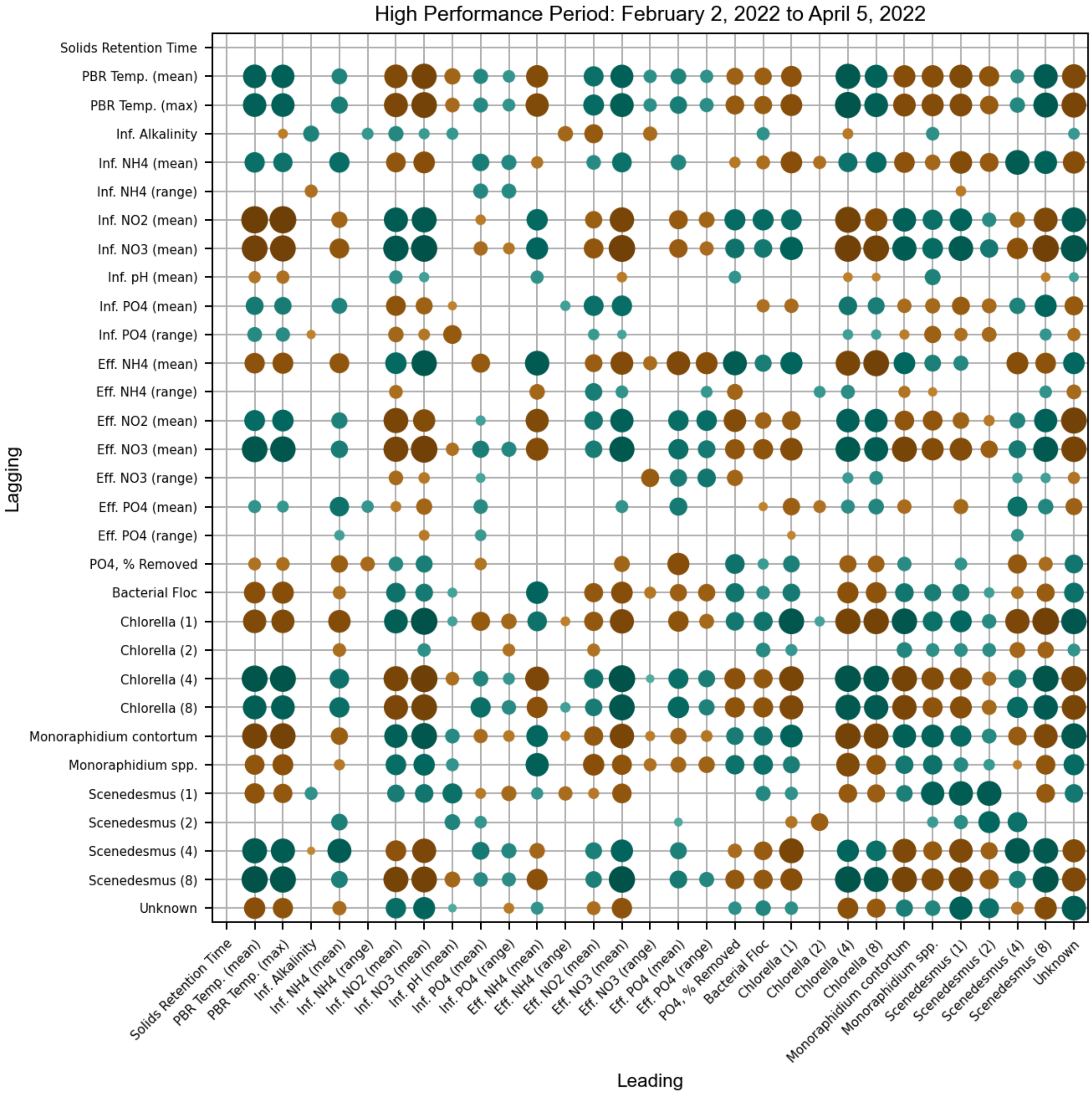


**Figure S6:** Asymmetric correlation matrix of key EcoRecover system parameters during a period of high performance (high orthophosphate recovery rate). In pairwise analysis, variables on the vertical axis have a time-lag with the variables on the horizontal axis. Mean and range quantities describe statistics on a per-sample-day basis (i.e., daily range). The view of microalgae biomass (**Figure S4**) selected for analysis is of total area. Spearman rank correlation coefficients are indicated by size and color: green = positive correlation, brown = negative correlation. Positive correlation describes time series that move in the same direction (e.g., both decreasing) and negative correlation describes movement in opposite directions (e.g., one increasing, other decreasing). Indices without statistical significance ($p < 0.05$) are omitted.

*Supplementary Figure S7:*

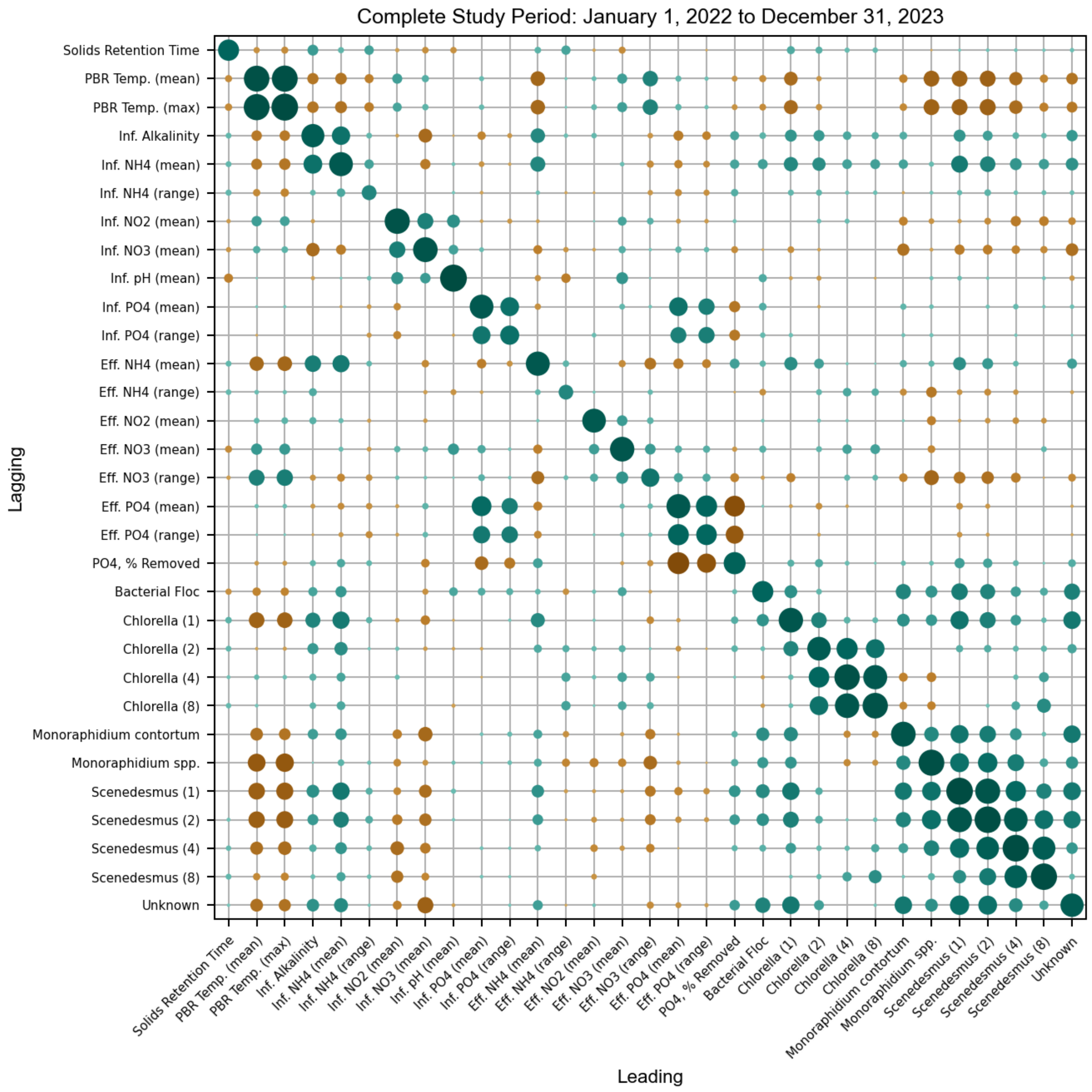


**Figure S7:** Asymmetric correlation matrix of key EcoRecover system parameters during the complete study period (variable orthophosphate recovery rate). In pairwise analysis, variables on the vertical axis have a time-lag with the variables on the horizontal axis. Mean and range quantities describe statistics on a per-sample-day basis (i.e., daily range). The view of microalgae biomass (**Figure S4**) selected for analysis is of total area. Spearman rank correlation coefficients are indicated by size and color: green = positive correlation, brown = negative correlation. Positive correlation describes time series that move in the same direction (e.g., both decreasing) and negative correlation describes movement in opposite directions (e.g., one increasing, other decreasing). Indices without statistical significance ($p < 0.05$) are omitted.
